# *NR1H3* p.Arg415Gln Is Not Associated To Multiple Sclerosis Risk

**DOI:** 10.1101/061366

**Authors:** The International Multiple Sclerosis Genetics Consortium, Chris Cotsapas

## Abstract

A recent study by Wang *et al* claims the low-frequency variant *NR1H3* p.Arg415Gln is pathological for multiple sclerosis and determines a patient’s likelihood of primary progressive disease. We sought to replicate this finding in the International MS Genetics Consortium (IMSGC) patient collection, which is 13-fold larger than the collection of Wang *et al*, but we find no evidence that this variant is associated either with MS or disease subtype. Wang *et al* also report a common variant association in the region, which we show captures the association the IMSGC reported in 2013. Therefore, we conclude that the reported low-frequency association is a false positive, likely generated by insufficient sample size. The claim of *NR1H3* mutations describing a Mendelian form of MS - of which no examples exist - can therefore not be substantiated by data.

---

In a recent study, Wang *et al* fail to demonstrate association to multiple sclerosis risk on chromosome 11p11.2 in the locus encoding *NR1H3* [1,2]. They sequenced the exomes of two affected individuals from a multiplex family (designated MS1) and selected 37 low-frequency missense variants found in both individuals for further study. After genotyping these variants in the nine members of that pedigree – including the initial two cases – and 185 controls, they excluded 33/37 variants and then genotyped the remaining four variants in 2,053 MS patients and 799 healthy controls. They detected one variant, rs61731956 (*NR1H3* p.Arg415Gln; GenBank: NM_005693.3; c.1244G>A), in a single additional case from another multiplex family (MS2), and from segregation analysis inferred that another four MS2 individuals were carriers, though only one of these has MS. Based on these four heterozygous cases, they claim linkage with a LOD score of 2.2, though the commonly accepted standard for significance is LOD > 3.0 [3].

We sought to validate the association of rs61731956 with MS susceptibility in our ongoing study of low-frequency missense variation in MS. After stringent quality control, we used linear mixed models to meta-analyze 32,852 cases and 36,538 controls of European ancestry in 14 country-level strata, genotyped for 250,000 low-frequency non-synonymous variants across all exons using Illumina’s HumanCore Exome array. We detected the minor allele rs61731956-A in nine of our strata, but find no evidence of association with overall MS risk (meta-analysis β = 0.06, p=0.32; Table 1). As Wang *et al* report this association specifically with PPMS, we compared 1,399 PPMS cases to 13,537 RRMS cases directly in six strata with available clinical course information, and also find no evidence of association with disease subtype (meta-analysis β = 2.35, p = 0.39; Table 2). Our previous linkage analysis of >700 multiplex families [4] further supports this conclusion (multipoint LOD = 0.0), as does earlier work from the Canadian Collaborative Project on the Genetic Susceptibility to Multiple Sclerosis (CCPGSMS) with no evidence of linkage in 40 Canadian families with four or more affected individuals, authored by members of the Wang *et al* study team [5].

**Table 1.** *NR1H3* p.Arg415Gln is not associated to multiple sclerosis risk. We metaanalyzed 32,852 cases and 36,538 controls of European ancestry in 14 country-level strata using linear mixed models to control for stratification. We detected the minor allele rs61731956- A in nine of our strata, but find no evidence of association to overall MS risk.

|  | Alleles |  | Minor allele frequency |  |  | Case allele counts |  | Control allele counts |  | p | Association |  |  |
| --- | --- | --- | --- | --- | --- | --- | --- | --- | --- | --- | --- | --- | --- |
|  | Minor | Major | All | Cases | Controls | A1 | A2 | A1 | A2 |  | OR | Cochran's Q | I<sup>2</sup> |
| Denmark | A | G | 0.00020 | 0 | 0.00040 | 0 | 2,534 | 1 | 2,475 | 0.375 |  |  |  |
| Germany | A | G | 0.00025 | 0 | 0.00026 | 2 | 8,950 | 3 | 11,425 | 0.845 |  |  |  |
| Italy | A | G | 0.00048 | 0.00065 | 0.00032 | 2 | 3,058 | 1 | 3,161 | 0.677 |  |  |  |
| Netherlands | A | G | 0.00042 | 0.00087 | 0.00028 | 1 | 1,151 | 1 | 3,609 | 0.317 |  |  |  |
| Norway | A | G | 0.00116 | 0.00127 | 0.00098 | 2 | 1,572 | 1 | 1,015 | 0.822 |  |  |  |
| Sweden | A | G | 0.00041 | 0.00053 | 0.00027 | 7 | 13,139 | 3 | 10,989 | 0.331 |  |  |  |
| UK/Australia | A | G | 0.00060 | 0.00054 | 0.00064 | 9 | 16,559 | 15 | 23,263 | 0.963 |  |  |  |
| USA/Boston | A | G | 0.00050 | 0.00042 | 0.00060 | 5 | 11,931 | 6 | 10,046 | 0.955 |  |  |  |
| USA/UCSF | A | G | 0.00048 | 0.00083 | 0.00000 | 3 | 3,627 | 0 | 2,660 | 0.127 |  |  |  |
| Meta-analysis | A | G |  |  |  | 31 |  | 31 |  | 0.3293 | 1.064 | 0.825 | 0 |

**Table 2.** *NR1H3* p.Arg415Gln is not associated to MS clinical course. We compared 983 PPMS cases and 7,72 RRMS cases directly in five strata with available clinical course information, and also find no evidence of association between rs61731956-A and clinical subtype.

| | Frequency in PPMS cases | Effect size ($\beta$) | Standard error | P | Cochran's Q | I<sup>2</sup> |
| --- | --- | --- | --- | --- | --- | --- |
| Italy | 0 | na | - | - |  |  |
| Netherlands | 0 | na | - | - |  |  |
| Norway | 0.0013 | -0.08 | 0.19 | 0.66 |  |  |
| Sweden | 0.0006 | 0.09 | 0.09 | 0.34 |  |  |
| UK/Australia | 0.0007 | 0.11 | 0.11 | 0.34 |  |  |
| USA/UCSF | 0.0012 | -0.08 | 0.17 | 0.65 |  |  |
| Meta-analysis | - | 2.32 |  | 0.39 | 0.69 | 0 |

Based on their interpretation of segregation patterns for rs61731956, Wang *et al* go on to genotype common variants in the NR1H3 locus in 2,053 MS patients and 799 healthy controls, but fail to detect any association with overall MS risk. They then report that four of their familial cases have a clinical course consistent with that of primary progressive MS (PPMS), and perform a secondary, stratified analysis of clinical course with the five tagging SNPs. They reduce their sample size to 420 PPMS and 1,287 RRMS patients for whom clinical course information was available, and describe an association between rs2279238 (OR = 1.35, p = 0.001) and PPMS, but not RRMS, risk. The IMSGC has already reported a disease risk association in this region [6] based on 14,498 MS cases and 24,091 controls to rs7120737, which is 420kb away and in moderate LD with rs2279238 (r^2^ = 0.62, D’ = 0.82 in the 1000 Genomes CEU panel). We also genotyped rs3824866, a perfect proxy for rs2279238 (r^2^ = 1, D’ =1 in the 1000 Genomes CEU panel), which shows modest association to MS risk (p = 2.1 × 10^−5^). Conditioning on rs7120737 fully explains this association, indicating that the result reported by Wang *et al* is a modest proxy for the strong signal we have previously reported.

Our 13-fold larger dataset therefore supports a more conventional interpretation of the data presented by Wang *et al*: there is no association between the low frequency *NR1H3* p.Arg415Gln variant rs61731956 and MS risk, but a common haplotype spanning the *NR1H3* locus is associated with overall MS susceptibility, despite the failure of Wang *et al* to detect it in their modestly sized cohort. Our data does not support an association specific to clinical course or PPMS. The false positive likely arose because Wang *et al* base their conclusions on a total of four affected carriers of the variant, and contravene standard practice by analyzing only five polymorphisms in the *NR1H3* locus, not controlling for population stratification, and failing to meet rigorous thresholds of significance for common variation (p < 5 × 10 ^−8^) or for family-based linkage (LOD > 3) [3,7].

Beyond these technical issues, Wang *et al* appear to have succumbed to an error in logic in their analysis. Although individually rare, coding variants are exceedingly common in the population: of 7,404,909 variants identified by the Exome Aggregation Consortium (ExAC) in 60,706 individuals, 99% have a minor allele frequency of <1% and 54% are seen exactly once in those data [8]. Therefore, there is complete certainty of observing at least one such variant in two closely related individuals, as Wang *et al* have done. This is reinforced by their observation of rs61731956-A in multiple unaffected individuals and the presence of this variant in 21/60,706 unselected ExAC individuals. This is not a unique false positive finding, as previous studies of equivalently small sample size have reported MS risk associations to low-frequency coding variants in *CYP27B1* [9] and *SAIE* [10], with both results failing to replicate in much larger studies with adequate statistical power [11–13].

We note that the experimental demonstration that p.Arg415Gln alters the heterodimerization efficiency between the NR1H3 product liver X receptor alpha and the retinoid X receptor alpha has no bearing on the association to MS pathogenesis. Many non-synonymous variants have dramatic effects on protein function, but in the absence of robust association to disease this alone cannot support a pathogenic argument [14].

The combination of our negative results from a 13-fold larger dataset and the methodological and logical flaws in the work presented by Wang *et al* categorically refute the bold claim that *NR1H3* variation defines a Mendelian subtype of MS, which would be the first monogenic form of the disease ever described. Such a discovery would have enormous implications for diagnosis of a subset of cases, prognosis and genetic counselling of extended family members, and eventually for clinical management of the disease in carriers. Unfortunately, the evidence provided by Wang *et al* does not support this conclusion.

